# The role of gene duplication in facilitating divergent patterns of gene expression across a complex life cycle

**DOI:** 10.1101/2024.01.30.577993

**Authors:** James G. DuBose, Jacobus C. de Roode

## Abstract

Explaining the processes that facilitate divergence in the morphologies and functions expressed by organisms throughout their life cycles is fundamental for understanding life cycle evolution. Theory suggests that the expression of traits is decoupled across life stages, thus allowing for evolutionary independence. Although trait decoupling between stages has been described in many studies, explanations of how said decoupling evolves have seldom been considered. Here, we propose evolutionary divergence between duplicate genes as an important mechanism by which life cycle complexity evolves. Because the different phenotypes expressed by organisms throughout their life cycles are coded by the same genome, trait decoupling between stages must be mediated through their divergence in gene expression. Gene duplication has been identified as an important mechanism that enables divergence in gene function and expression between cells and tissues. Here, we examined the temporal changes in gene expression across the monarch butterfly (*Danaus plexippus*) metamorphosis. We found that within homologous groups, more phylogenetic divergent genes exhibited more distinct temporal expression patterns, and that this relationship scaled such that more phylogenetically diverse homologous groups showed more diverse patterns of gene expression. Furthermore, we found that duplicate genes showed increased stage-specificity relative to singleton genes. Overall, our findings suggest an important role of gene duplication in the evolution of trait decoupling across complex life cycles.

**Significance:** The proliferation of many of the world’s most diverse groups of eukaryotes is frequently attributed to their life cycle complexity. By allowing organisms to express different traits throughout their lives, complex life cycles enable individuals to utilize multiple ecological niches. However, the mechanisms that facilitate life cycle evolution are not well understood. We drew inspiration from studies on functional divergence between different tissues and examined the role of gene duplication in generating different patterns of gene expression between stages across the metamorphosis of *Danaus plexippus* (the monarch butterfly). Our findings suggest that the role of gene duplication in generating differences between cell and tissue types likely extends to trait differentiation between stages within complex life cycles.

## Introduction

Many groups of organisms undergo extensive morphological and ecological shifts throughout their life cycles. These shifts appear gradual in some organisms, as seen in the relatively continuous development from infant to adult in primates. However, these shifts seem more complex in many other organisms; a larva first must transition into an intermediate pupal stage before restructuring its morphology into the form of a butterfly. Changes in life cycle complexity have been associated with the diversification of many taxa throughout natural history (1, 2). Despite nearly a century of interest in the evolution of complex life cycles, we still lack a general understanding of the mechanisms that facilitate divergence in the morphologies and functions expressed by organisms throughout their lives.

This gap in our understanding can be partially attributed to the view that complex life cycles are divided into stages that are discrete, which is the central assumption made in foundational theoretical work (3, 4). This simplifying assumption limits existing hypotheses, as it is apparent that the transition from one life stage to the next requires continuous changes in the relative abundance, activity, or placement of different cells and tissues (5). Therefore, we propose that life cycle evolution can be more fundamentally described by the body of theory concerning the evolution of cell and tissue differentiation, which focuses on describing the mechanisms that facilitate evolutionary change in gene function and continuous patterns of expression.

The adaptive decoupling hypothesis is the most prominent explanation for the evolution of complex life cycles, and is an extension of an earlier hypothesis that different life stages adapt independently to the niches they occupy (3, 4). The adaptive decoupling hypothesis elaborates that complex life cycles allow different stages to independently respond to natural selection by genetically decoupling the development of their traits (3). This hypothesis predicts that genetic variation should generate phenotypic variation in certain life stages but not others. Many studies have found results consistent with this prediction (6–9), and more recent studies have elucidated variation in gene expression between stages as the likely driver of said genetic independence (10, 11). However, the predictions made by the adaptive decoupling hypothesis are limited to descriptions of extant signatures of decoupled traits, which fails to provide a mechanism that explains how trait decoupling evolves. We propose that the evolution of trait decoupling can be explained by the mechanisms already established in the evolution of cell and tissue differentiation because transitions between life stages are driven by continuous turnover in cell and tissue types.

Gene duplication is the best described mechanism that generates evolutionary change in patterns of gene expression between cells and tissues (12–15), though other mechanisms have been less studied (16–18). Gene duplication can be generated though unequal crossing over, retrotransposition, and chromosomal duplication and provides a rich source of genetic variation that can facilitate major evolutionary change (19, 20). Hypotheses concerning the evolution of duplicate genes share key similarities with the adaptive decoupling hypothesis. For example, the neofunctionalization hypothesis suggests that retention of the ancestral function in one copy alleviates selective constraints on the other copy, allowing it to develop novel functions that are more subject to selection (19). However, a role of neutral evolution in generating functional divergence between duplicate genes has also been described (12, 21), thus offering a more comprehensive mechanism by which traits could diverge between stages. More generally, the idea that complexity is added to a genome through gene duplication is well established (19, 22, 23), and empirical evidence for gene duplication resulting in more complex phenotypes has been documented in a variety of taxa (24–27). Therefore, investigating the role of gene duplication in life cycle evolution has potential to broaden our understanding of how biological complexity evolves.

Although there are several nuances to predicting the relationship between sequence evolution and expression pattern evolution in duplicate genes, the general expectation is that the evolution of duplicate genes leads to more divergent and (stage) specific expression patterns (13, 14). While this insight has primarily been derived from relating duplicate gene evolution to expression pattern divergence between different mammalian tissues, we hypothesize that the same patterns will emerge when examining temporal patterns of gene expression across a complex life cycle. To test the predictions that the evolution of duplicate genes leads to more divergent and stage-specific expression patterns (Figure 1), we examined patterns of gene expression across the holometabolous life cycle of the monarch butterfly, *Danaus plexippus.* The *D. plexippus* life cycle is characterized by a non-dispersive caterpillar stage that is specialized for feeding on milkweed foliage, followed by a non-feeding pupal stage during which metamorphosis occurs, and a final highly dispersive imaginal (butterfly) stage that is specialized for reproduction and feeding on nectar. The extreme ontogenetic niche shifts and trait divergence between stages makes *D. plexippus* a promising model system for studying the evolutionary processes that generate morphological and functional divergence throughout life cycles.

**Figure 1.**
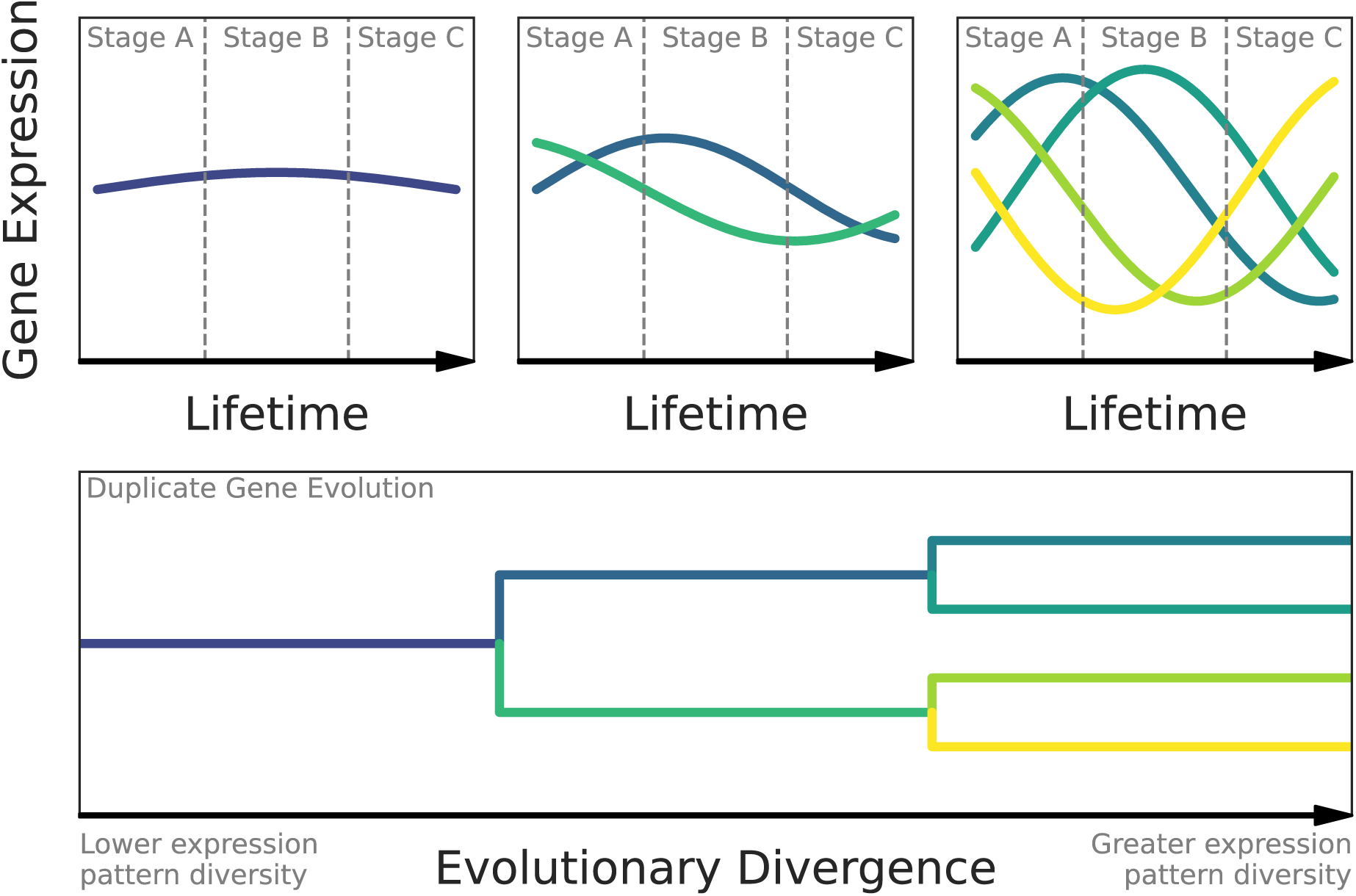
A conceptual diagram showing the hypothesized mechanism of how duplicate gene evolution could lead to divergence in gene expression (and consequently phenotypes) between perceived stages in a complex life cycle. Initially, a given gene has an expression pattern that is relatively uniform throughout an organism’s lifetime. After duplication, the expression patterns of each copy tend to diverge and become more stage specific. After additional duplication and divergence, expression tends to diverge and specify even more between copies. This makes the expression at each stage substantially more distinct from other stages, which would result in greater phenotypic divergence between stages if the duplicates functionally diverged as well. This diagram does not show all possible fates of duplicate genes.

## Methods

### Experimental design and D. plexippus rearing

To quantify changes in gene expression across the holometabolous development of *D. plexippus*, we sequenced mRNA extracted from third instars, fifth instars, early pupae (one day after pupation), late pupae (6-8 days after pupation), and adults (several hours after eclosion). A previous study has suggested that feeding on more toxic milkweed induces changes in gene expression during the second instar (28). Therefore, we reared larvae on both *Asclepias incarnata* (less toxic) and *Asclepias curassavica* (more toxic) to ensure that our findings are robust to a major source of environmental variation. We collected five individuals at each stage and from each plant for mRNA quantification.

Parental (P) *D. plexippus* butterflies were caught in St. Marks, Florida, U.S.A. (30°09′33″N 84°12′26″W) during October of 2022. Butterflies were overwintered in a 14°C incubator (to maintain a state of diapause) and were fed approximately 10%-20% honey water every ten days. During March of 2023, butterflies were mated to establish an F1 generation. F1 caterpillars were reared on *Asclepias curassavica* and after maturation and mating, the F2 caterpillars used in this experiment were reared on either *A. curassavica* or *A. incarnata.* To reach the necessary sample size, we used F2 caterpillars from two different lineages that did not share P or F1 ancestors. Treatments of plant species and development stage were randomly distributed to caterpillars from both lineages to minimize confounding due to genetic background. All individuals sampled in this study were reared at the same time and in the same conditions (See Appendix section 1.1-1.3 for details).

### Sample collection and preparation

To minimize possible effects of sample handling, all caterpillars, pupae, and adults were snap frozen in liquid nitrogen before being stored at -80°C. Third instars, fifth instars, early pupae, and late pupae were all frozen in sterile centrifuge tubes, and adults were frozen in glassine envelopes several hours after eclosion (after their wings had finished expanding). For each day freezing took place, samples were stored in a polystyrene foam cooler full of dry ice until all flash freezing for that day was completed. This process took approximately one hour or less on any given day, so no sample was on dry ice for more than an hour before being transferred to the -80°C freezer. All freezing took place in the same greenhouse room that the caterpillars were reared in, and no individual left said room before being frozen throughout the duration of the experiment.

Because we were interested in global gene expression patterns, we collected samples by homogenizing whole bodies using a sterile porcelain mortar and pestle. Each sample for a given round of homogenization was placed in a cooler filled with dry ice. Samples were individually placed in a mortar and liquid nitrogen was constantly added throughout the homogenization to prevent samples from thawing. After a given sample was completely homogenized, homogenate was quickly collected using a sterile polypropylene spatula and stored in a fresh centrifuge tube. Twenty samples were randomly selected for each round of homogenization.

### RNA extraction and sequencing

We used a Promega SV Total Isolation System kit to extract total RNA from *D. plexippus* homogenate. While our workflow generally followed the manufacturer’s suggested protocol, we made several alterations to obtain higher quality RNA. Briefly, we doubled the recommended RNA lysis buffer to increase the relative amount of RNA at the initial lysis step. We also added an additional centrifugation step after the initial tissue lysis to further clear organic contaminants and improve final extract purity. All centrifugation steps were increased to 20,000 rcf to better clear organic contaminants and performed at 17°C to avoid sample heating. Each batch of extractions consisted of 11 randomly selected samples and 1 negative control. After each extraction, we used a NanoDrop to quantify the purity and concentration of the RNA. Samples with an A260/A280 or an A260/A230 of less than 1.95 were discarded and re-extracted.

After all extractions were completed, purified RNA was packaged in dry ice and sent to Novogene (Sacramento, CA) for library preparation and sequencing. Briefly, Novogene used an Agilent 5400 Fragment Analyzer System to confirm that all samples had adequate purity levels, concentrations, and volumes, as well as acceptable RNA integrity numbers (minimum = 7.9). Libraries were then prepared via poly-A tail selection and sequenced using a 150bp paired-end approach on a NovaSeq 6000 sequencing system, thus ensuring at least 20 million reads were obtained for each sample.

### Sequence processing and gene expression quantification

Initial quality control of raw sequences was performed by Novogene, where adapter sequences, reads with ambiguous base calls in greater than 10% of the read, and reads with a phred score of less than or equal to 5 in 50% of the read were removed. After receiving the sequences from Novogene, we used FASTQC to generate additional quality reports for each sample (29). This showed that the median phred score did not drop below 30 at any position for any sample. Therefore, no additional sequence quality control was performed.

To quantify transcript abundances for each gene, we used kallisto (v.0.46.2) to pseudo-align reads to the coding sequences of the *D. plexippus* reference genome (v.Dpv3, GenBank Assembly = GCA_000235995.2) (30). Downstream analyses were performed using transcript per million normalized read counts (automatically generated by kallisto) to minimize biases due to unequal gene lengths and varying library sizes (31, 32).

### Quantifying gene expression divergence between stages

Given the high dimensionality of gene expression data, we first computed the Manhattan distance between each sample using the *dist* R function (33). We then used the *adonis2* function from the *vegan* R package (v.2.6-4) (34) to perform a permutational multivariate analysis of variance (PERMANOVA) with 999 permutations, where developmental stage and plant were initially considered as factors. We then performed a PERMANOVA on each set of adjacent stages, as well as between each larval stage and the adult stage. To visualize global expression divergence between stages, we performed principal coordinate analysis using the *prcomp* R function (33).

### Quantifying the relationship between gene phylogenetic divergence and expression pattern divergence within homologous groups

To infer homology between genes, we first used PSI-BLAST (BLAST 2.5.0+) (35) with five iterations to align all *D. plexippus* protein sequences to each other. Genes were then inferred to be homologous if the query sequence showed at least 30% similarity across the length of the target sequence, as well as an E-value of at least 1x10^-10^. To examine how including more distant homologs could impact our analysis, we performed an additional analysis where homology was inferred based on at least 20% similarity across 70% of the target sequence and an E-value of less at least 1x10^-5^. These less stringent similarly cutoffs for homology inference showed consistent results with our primary analysis (Appendix section 3.2). Homologous pairs were assembled into sets of two-node subgraphs, and subgraphs were then merged based on common node identity to assemble homologous groups.

To quantify the phylogenetic distance between members of inferred homologous groups, we first used MUSCLE (v.5.1) to create a multiple sequence alignment for each group (36). We then used IQ-TREE2 (v.2.1.4) to identify the best fit sequence evolution model and infer maximum likelihood phylogenies for each multiple sequence alignment (37, 38). We then used the *cophenetic.phylo* function from the *ape* R package (v. 5.7-1) (39) to calculate pairwise phylogenetic distances from each homologous group tree. To calculate pairwise expression pattern distances, we first mean centered and standardized the median transcripts/million value for each gene within each stage to better measure distance between temporal patterns as opposed to magnitude (which cannot be assessed with our data). We then calculated the pairwise Euclidian distance between each gene expression pattern within a given homologous group using the *dist* R function (33). Finally, we used Mantel tests to calculate the correlation between phylogenetic and expression pattern distance matrices for each homologous group, which were implemented via the *mantel* function in the *vegan* R package (v.2.6-4) (34). We then used a t-test to test if the distribution of correlation coefficients was positively shifted from 0, which was implemented using the *t.test* R function (33).

### Quantifying the relationship between phylogenetic diversity and expression pattern diversity across homologous groups

The diversity (*D*) of each previously described phylogenetic tree was calculated as the sum of branch lengths: *D* = ∑^n^_i = 1_ *l_j_*, where *n* represents the number of branches and *l_i_* represents the length of the *i*th branch. To quantify expression pattern diversity, we first used the Ward method to created hierarchical clustering graphs of the temporal expression patterns for each gene. Prior to clustering, the transcripts/million values for each gene were mean centered and standardized because hierarchical clustering will group expression patterns that show distinct temporal trends but have more similar average relative abundances across time points. For each hierarchical clustering graph, diversity was calculated as previously described for phylogenetic diversity. We then fit a linear model to examine the relationship between phylogenetic diversity and expression pattern diversity across all inferred homologous groups. Because diversity was calculated additively (for each branch, diversity was added in proportion to divergence), we also fit individual linear models to each homologous group size that had at least five replicates. In addition to removing the inherent positive correlation between group size and diversity, this approach also allowed us to compare global and local patterns. All hierarchical clustering graphs were constructed using the *hclust* R function and all linear models were fit using the *lm* R function (33).

### Null model algorithm and analysis

To test if observed patterns could be randomly derived from our data we constructed a null model that assigns a randomly sampled expression pattern to each gene for each homologous group and recalculates the linear relationship between phylogenetic and expression pattern diversity:

Consider the total pool of observed expression patterns *E = {X_1_, X_2_, X_3_,…,X_i_}*, where *X* represents an expression pattern and *i* represents the number of genes in the *D. plexippus* genome. Previously defined groups of homologs are represented as *H = {G_1_, G_2_, G_3_,…,G_j_}*, where *j* represents the number of homologous groups and also serves as the index for the set of observed phylogenetic diversities for each group, represented as *P = {D_1_, D_2_, D_3_,…,D_j_}.* For each group *G_j_*, *G_j_ = {X_i_1, X_i_2, X_i_3,…,X_i_n}*, where *n* represents the number of genes included in each group and *X_i_n* represents a corresponding expression pattern from *E*. Note that not all expression patterns in *E* are included in some *G_j_*. For each *X_i_n* in each *G_j_*, *X_i_n* ← *X_i_k*, where *X_i_k* represents the *kth* expression pattern randomly selected with replacement from *E* for the *jth* group. After random assignments of expression patterns, expression pattern diversity *S_j_* is calculated for each *G_j_* as previously described, which is represented as *R = {S_1_, S_2_, S_3_,…,S_j_}. R* is then combined with *P* to produce *N* such that *N = {(D_j_, S_j_) | D_j_ ∈ P, S_j_ ∈ R}*. Finally, a linear model is then fit both globally and to each group size, as previously described for the observed analysis.

This algorithm was repeated 1000 times to generate null distributions of global and group-size specific linear relationships between phylogenetic and expression pattern diversity. To evaluate the observed results against the null model predictions, we calculated the probability that the null model would produce an effect greater than the observed, as well as the standardized effect sizes 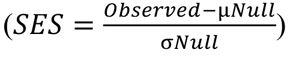 for the slope of each linear model.

### Expression specificity calculation and analysis

Stage-specificity for each gene was calculated using the tissue specificity index τ (40), which ranges from 0 (equal expression across stages) to 1 (expression in a single stage): τ = 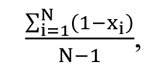 where *N* is the number of stages (for our purposes) and *x_i_* is the expression level 6,$ normalized to the maximum expression value across stages. Although τ was developed for assessing tissue specificity, it has been used to gain insight into temporal specificity as well (41). We then performed a Kolmogorov–Smirnov test using the *ks.test* R function (33) to assess if the distribution of τ values was shifted in duplicated genes relative to singleton genes.

## Results

### The extent of transcriptional divergence between D. plexippus larvae and pupae is comparable to the divergence between larvae and adults

Because all distinct phenotypes expressed throughout a complex life cycle are coded by the same genome, trait decoupling must be mediated through variation in gene expression across stages. Therefore, we were first interested in the extent that gene expression has diverged between stages throughout the *D. plexippus* metamorphosis.

Overall, we found that gene expression significantly varied by developmental stage (F = 61.36, p < 0.001) but not plant host (F = 0.88, p = 0.47) (Figure 2). We then performed pairwise comparisons to test for differences between subsequent stages, as well as between larvae and adults. Following *D. plexippus* throughout metamorphosis: the transition from third instar to fifth instar involves some, but relative few changes in gene expression (distance = 6.97x10^5^, F = 18.67, p < 0.001). Then a substantial change in gene expression occurs during the transition from fifth instar to early pupa (distance = 1.22x10^6^, F = 68.06, p < 0.001), followed by a slightly smaller but comparable change from early pupa to late pupa (distance = 1.20x10^6^, F = 62.25, p < 0.001). Finally, the transition from late pupa to adult involves a modest change in gene expression (distances = 9.37x10^5^, F = 35.43, p < 0.001), but said change is notably less than the changes involved in the previous two transitions. It’s interesting to note that the extent of divergence in gene expression between fifth instars and early pupae is comparable to the divergence between both larval stages and adults (third instar: distance = 1.16 x10^6^, F = 108.08, p < 0.001; fifth instar: distance = 1.24 x10^6^, F = 71.65, p < 0.001). This distinction in early pupae appears to involve a decrease in metabolic investment and an increase in immune investment (Appendix Figure 6). More broadly, the transcriptional changes across stages appear to be mostly driven by differential investment in metabolism and genetic information processing, consistent with niche shifting and developmental requirements (Appendix Figure 6).

**Figure 2.**
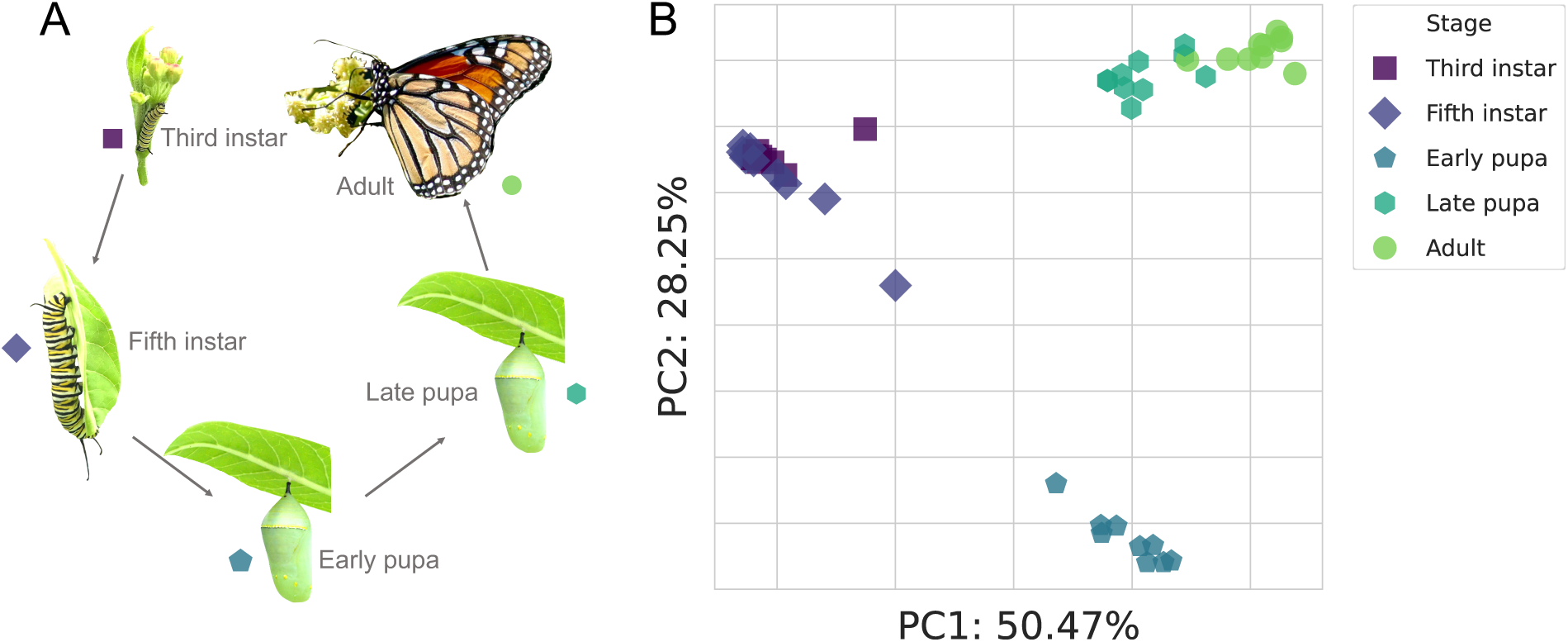
A depiction of how morphology and transcription changes across the *D. plexippus* lifecycle. A) Images of each life stage sampled in this study showing. B) A principal coordinate analysis plot showing substantial transcriptional divergence between life stages. Each point represents the global gene expression profile of an individual, and closer points indicate more similar gene expression profiles. Axis labels indicate principal coordinate rank and the proportion of variance explained.

### Phylogenetic divergence between homologs generally corresponds with increased divergence in temporal expression pattern

As previously described, the general hypothesized outcome of evolutionary divergence between homologs is increased divergence in their expression patterns. Consistent with this hypothesis, we generally found a positive relationship between phylogenetic distance and expression pattern distance within homologous groups (Figure 3). Specifically, a positive association was observed in approximately 72% of groups. However, we note that there is variation in the both the strength and direction of said correlations, with the remaining 28% of groups showing null or negative correlations. Nonetheless, the distribution of correlation coefficients is shifted positively from 0 (mean = 0.19, t = 6.22, p = 7.23 x 10^-9^).

**Figure 3.**
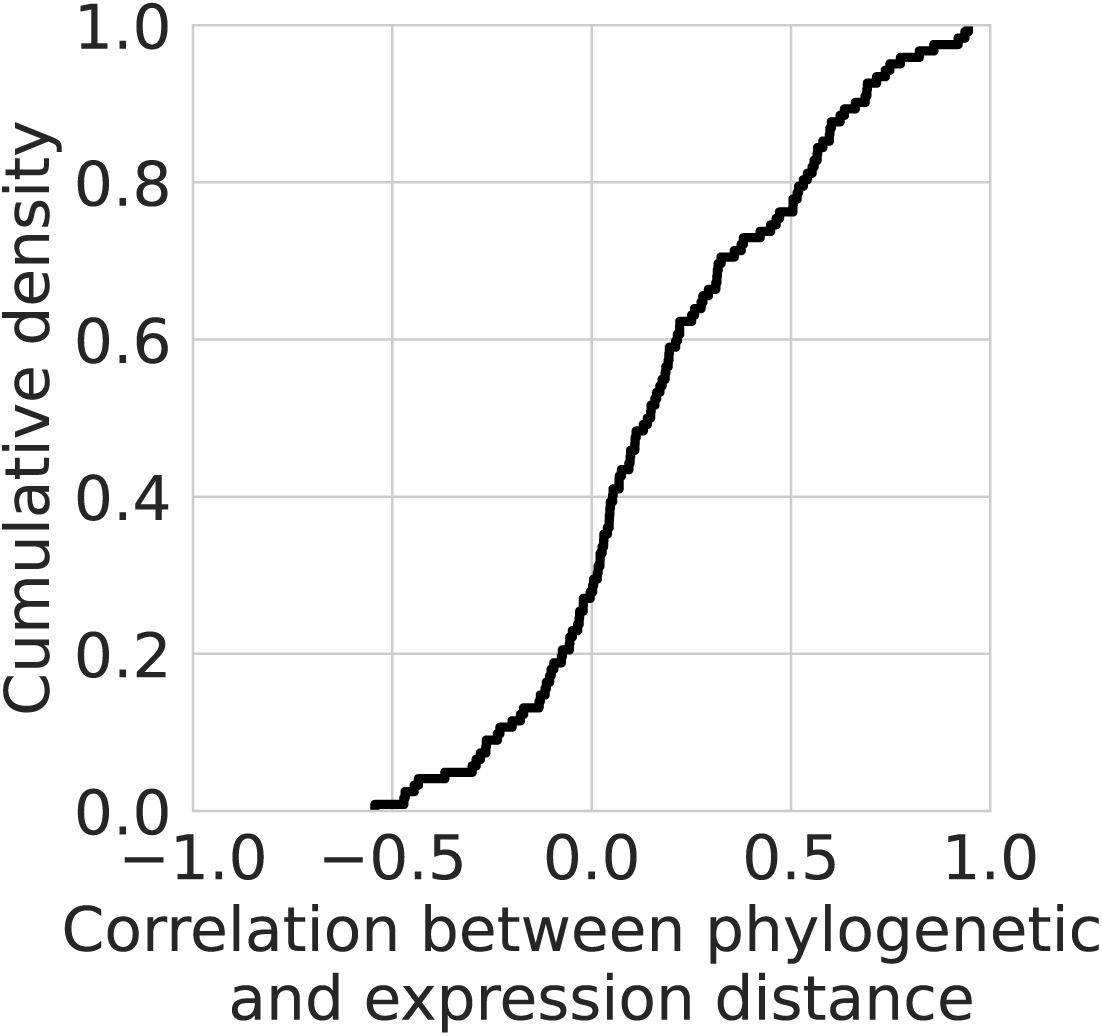
Phylogenetic distance positively correlates with expression pattern distance in most homologous gene groups. The empirical cumulative density function of correlation coefficients between phylogenetic distance and expression pattern distance. Values greater than 0 indicate a positive correlation and greater values indicate stronger correlations. Overall, the majority of the distribution (approximately 72%) consists of positive correlations.

### Diversity in the temporal expression patterns exhibited by homologous groups increases with their phylogenetic diversity

If expression pattern diverges with phylogenetic divergence between genes within a homologous group, the pattern that should emerge is an association between overall phylogenetic diversity and expression pattern diversity across homologous groups. Consistent with this hypothesis, we found a positive relationship between phylogenetic diversity and expression pattern diversity (Figure 4). However, we also found that the increase in expression pattern diversity started to saturate at higher phylogenetic diversities, and that this relationship was better described by a quadratic model than a linear model (linear model SSE = 780.55, quadratic model SSE = 750.61).

**Figure 4.**
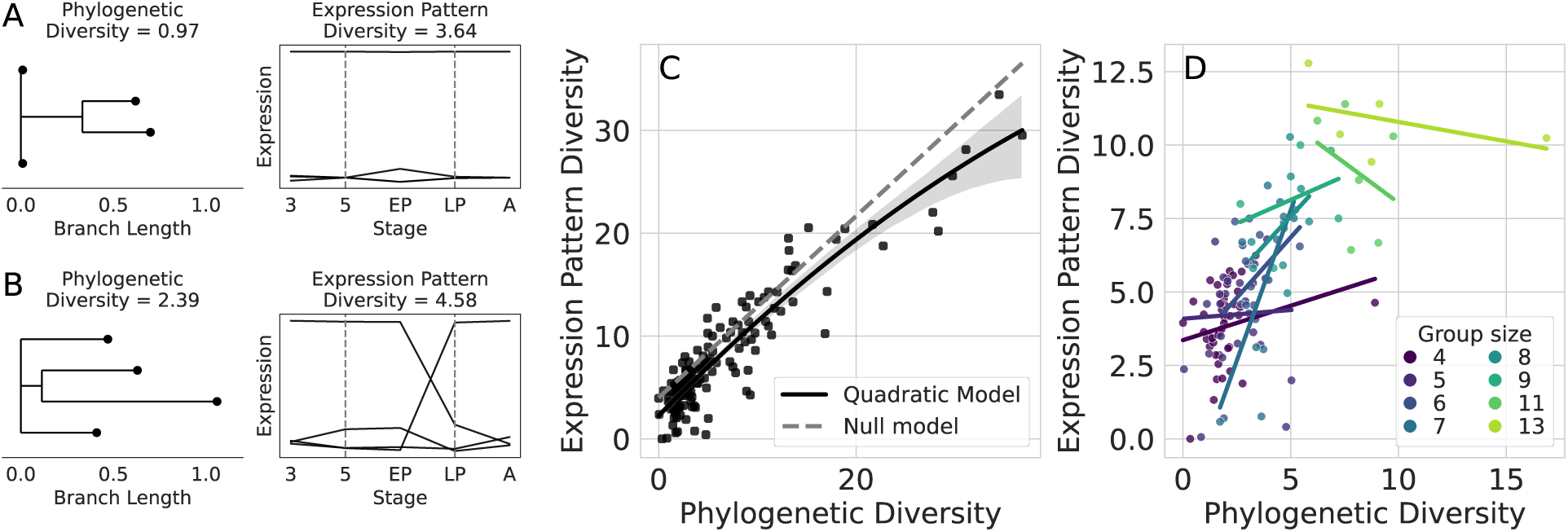
More phylogenetically diverse homologous groups exhibit more diverse patterns of expression. A) An example of a less phylogenetically diverse homologous group (a geranylgeranyl diphosphate synthase-like group) showing less diverse patterns of expression. B) An example of a more phylogenetically diverse group (an arrestin-like group) showing more diverse patterns of expression. A and B are meant to contextualize the broader analysis, not to lend interpretation about the specific homologous groups used for demonstration purposes. C) The relationship between phylogenetic and expression pattern diversity across all homologous groups. The solid black line depicts the fit quadratic model, the light gray area indicates the 95% confidence interval for said model, and the dashed line shows the average null model prediction. D) The relationship between phylogenetic and expression pattern diversity by homologous group size. Each line represents the linear model fit to each homologous group size (only group sizes with 5 or more replicates were considered in this analysis).

The positive relationship between phylogenetic and expression pattern diversity could have been driven by the addition of genes to homologous groups, as opposed to phylogenetic diversification within the group. Therefore, we examined the relationship within each homologous group size that showed sufficient replicates for fitting a linear model. This analysis revealed positive correlations between phylogenetic and expression pattern diversification for each of the six smaller homologous group sizes (mean correlation coefficient = 0.35, range = [0.04, 0.68]), but negative associations in the two larger group sizes (mean correlation coefficient = -0.38, range = [-0.44, -0.31]).

The previous two analyses indicate a positive but saturating relationship between phylogenetic and expression pattern diversity. To test if this pattern could be randomly derived from our data, we constructed a null model that randomly assigns observed expression patterns to genes within homologous groups and examined the slopes of the relationships between phylogenetic and expression pattern diversity. Across all homologous groups, the null model consistently predicted that expression pattern diversity would increase at a higher rate with respect to phylogenetic diversity (SES = -4.07, *P(Observed > Null)* < 0.001) (Figure 4C). Consistent with the previous analysis, we found that the expression pattern diversity increased at a higher rate than predicted by the null model in each of the six smaller groups (mean SES = 2.34, 95% CI = [0.75, 3.94]) (Appendix Figure 3). Furthermore, the null model seldom predicted a rate of increase greater than or equal to the observed rate (*P(Null ≥ Observed)* ≤ 0.025 in four of the six smaller groups). In contrast, the two larger groups did not show significant deviation from null predictions (mean SES = -1.1*, P(Null ≥ Observed)* ≥ 0.81) (Figures 4B and 4C). Full results for observed-null comparisons can be found in Appendix Table 3.

### Genes within duplicated genes tend to show more stage-specific expression patterns than singleton genes

Another key prediction regarding expression pattern divergence between duplicate genes is that copies will show increased stage specificity. Consistent with this prediction, we found that genes within homologous groups tended to show increased stage specificity relative to singleton genes (D = 0.175, p < 2.2x10^-16^) (Figure 5). Furthermore, we found a weak but significant positive relationship between expression specificity and homologous group size (r = 0.108, p < 2.2x10^-16^). However, homologous group size explained very little variance in expression specificity (R^2^ = 0.0117).

**Figure 5.**
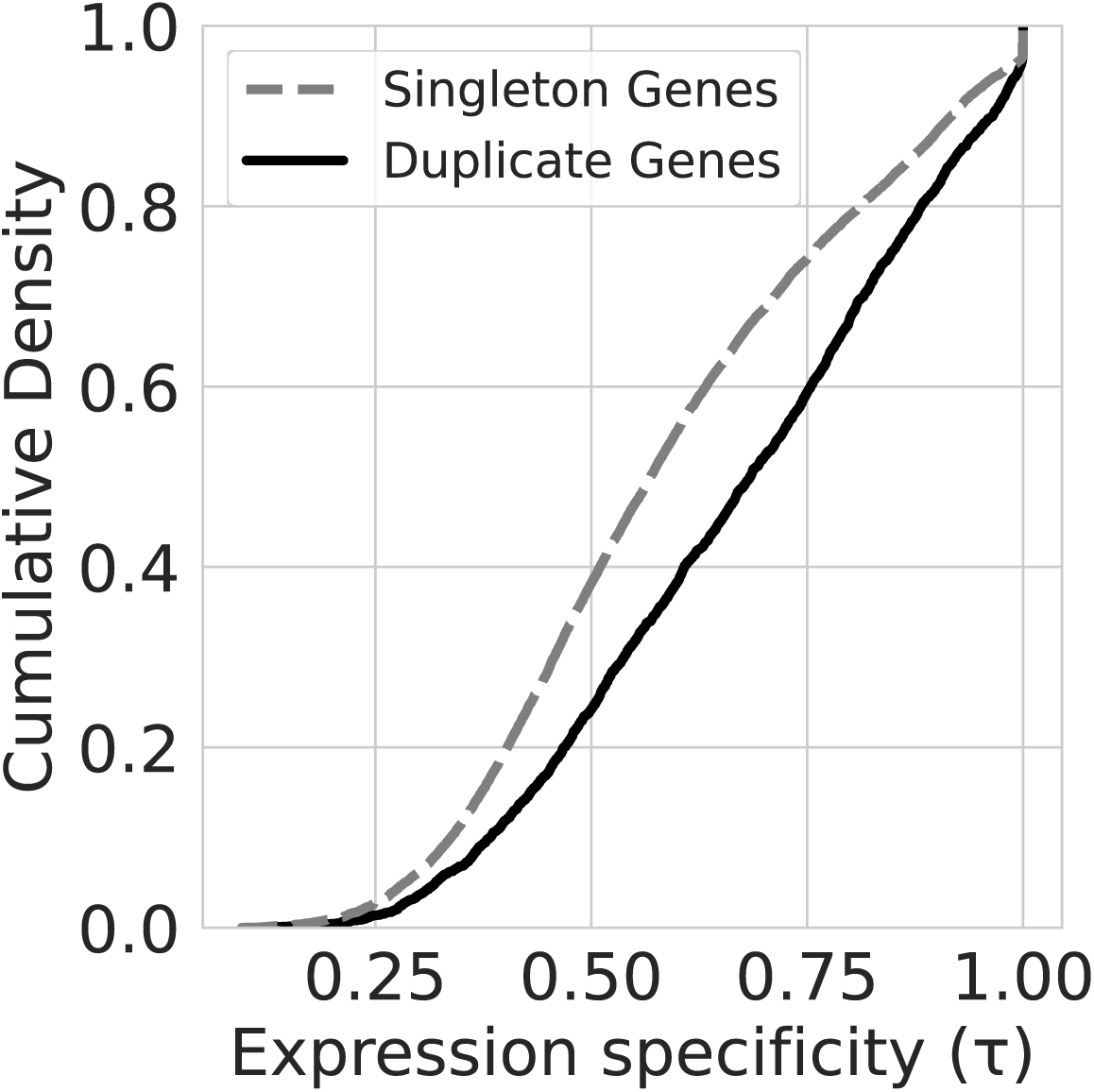
The expression patterns of duplicate genes show increased stage-specificity relative to singleton genes. The empirical cumulative density functions of expression specificity values for duplicate (solid line) and singleton (dashed line) genes. Higher expression specificity values indicate increased stage-specificity.

## Discussion

We found that phylogenetic divergence between genes generally resulted in more divergent patterns of gene expression within homologous groups (Figure 3), and that more phylogenetically diverse groups generally exhibited more diverse patterns of gene expression (Figure 4). Furthermore, we found that genes within homologous groups showed increased stage-specificity relative to singleton genes (Figure 5). As predicted, these results are consistent with studies that have examined the role of gene duplication in facilitating expression divergence between different cells and tissues (12–15, 40, 41). This consistency suggests that theories of evolution by gene duplication can be applied more generally towards explaining functional differentiation between stages at the organismal level.

Our findings significantly expand on previous findings that duplicate genes were more likely to vary in expression between larvae and pre-pupae in several *Drosophila* species than singleton genes. (42). A more nuanced pattern that we observed was a saturating relationship between phylogenetic diversity and expression pattern diversity, which could not be explained by our null model (Figure 4). This pattern was recapitulated across homologous group sizes, where the positive relationship between expression pattern diversity and phylogenetic diversity disappeared at larger and more diverse groups (Figure 4). Similar patterns have been documented in humans, mice, and yeasts, with expression divergence occurring more rapidly at shorter evolutionary time scales before plateauing at longer time-scales (12, 15, 43). Possible explanations for this pattern include decoupled rates of evolution in coding sequence and regulatory elements, dosage sensitivities/balancing, and additional complexities related to neo/sub-functionalization dynamics (18, 44–46). Regardless of the specific mechanisms, which are beyond the focus of this study, finding this consistency provides stronger evidence that our results recapitulate more fundamental work on duplicate gene evolution.

While our findings showed general agreement with previous studies, deviation from the predicted relationship between phylogenetic divergence and expression pattern divergence were also found. These deviations could likely be explained by the historical context in which specific homologous groups originated and evolved. For example, whether or not the duplication event was lineage-specific or occurred ancestrally, whether or not duplicates arose from a small-scale duplication event or a chromosomal duplication event, and relative importance of selective and neutral processes in generating sequence divergence, are all expected influence duplicate functionalization and expression divergence (12, 14, 43). Furthermore, gene duplication is not the only mechanism that facilitates evolutionary change in gene expression patterns. Genes are typically expressed as part of complex regulatory networks, and evolutionary changes to said networks can occur through other mechanisms not inherently related to gene duplication (17, 18). These mechanisms are not exclusive, and future studies that quantify the relative importance of duplicate gene evolution and regulatory network evolution in generating temporal divergence in gene expression will give key insight into the evolution of life cycle complexity.

We interpret our results as evidence for an important role of gene duplication in facilitating life cycle evolution. However, it is important to emphasize that we do not suggest that expansion of the specific homologous groups identified in our analyses were directly involved in the origin of holometabolous development; the origin of holometabolous development was not the focus of this study. Rather, our aim was to identify a general process by which traits become temporally decoupled, which would result in complex life cycles when said traits are accumulated over time. Previous studies have documented the patterns that emerge from temporal trait decoupling. Genetic independence of traits expressed by different stages has been well described (see: (6, 7, 47, 48) for examples and (49) for a detailed review), and more recent studies have elucidated variation in gene expression between stages as the likely cause of said independence (8, 10, 11, 50). Our findings are consistent with this interpretation as well. However, a common theme across previous studies is that decoupling is variable and not universal to all traits or genes. Therefore, a more mechanistic understanding of how decoupling evolves is needed to understand life cycle evolution more comprehensively. Our findings provide initial evidence for an important role of gene duplication in the decoupling of traits and more generally in facilitating divergence in temporal gene expression patterns across stages.

Because our samples consisted of whole bodies, the variation in gene expression observed between stages likely represents shifts in the relative abundance or activity of different cell and tissue types throughout the *D. plexippus* post-embryonic development. This, paired with the consistency of our findings with work on the role of gene duplication in generating functional differentiation between cells and tissues suggests that life cycle evolution in multicellular organisms can be more fundamentally understood through evolutionary shifts in the timing at which different cell and tissue types are expressed, which echoes Haldane’s earlier ideas (5). Therefore, we hypothesize that what is perceived as distinct stages could be explained by the extent of turnover in cell and tissue types within an organism. Likewise, what is perceived as discrete transitions between stages could be explained by the rate at which said turnover occurs. From this perspective, the continuous transition from infant to adult in primates could be mechanistically linked to the extreme transition from larva to butterfly in lepidopterans.

## Acknowledgements

We thank Christopher P. Catano and Mackenzie Hoogshagen for helpful comments and discussion on this work. We thank Erik Edwards for growing the plants used for *D. plexippus* rearing, and the members of the de Roode lab for help in managing monarch mating. This work was supported by National Science Foundation grant IOS-1922720 to J.C.dR.

## Author Contributions

J.G.D and J.C.dR designed and performed research. J.G.D analyzed the data. J.G.D and J.C.dR wrote the paper.

## Data and code availability

All sequences and count matrices generated for this project have been deposited in the NCBI GEO database and can be accessed with the accession number GSE253389 or the BioProject accession number PRJNA1065445. All code written for data analysis can be accessed at https://github.com/gabe-dubose/mtstp.

## Appendix

### 1 Extended Methods

#### 1.1 Study system and experimental design

Holometabolous development involves the transition from a larval stage that is typically specialized for feeding and growth to a stationary or less mobile pupal stage. During the pupal stage, dramatic morphological restructuring occurs, resulting in a distinct adult stage that is typically specialized for dispersal and reproduction.

To quantify changes in gene expression the across the holometabolous development of *Danaus plexippus*, we sequenced mRNA extracted from third instars, fifth instars, early pupae (one day after pupation), late pupae (6-8 days after pupation), and adults (several hours after eclosion). A previous study has suggested that feeding on more toxic milkweed induces changes in gene expression during the second instar [1]. Therefore, we reared larvae on both *Asclepias incarnata* (less toxic) and *Asclepias curassavica* (more toxic) to ensure that our findings are robust to a major source of environmental variation. We collected five individuals at each stage and from each plant for mRNA quantification. All individuals sampled in this study were reared at the same time and in the same conditions.

#### 1.2 Milkweed cultivation

*A. incarnata* and *A. curassavica* seeds were purchased from Joyful Butterfly (Blackstock, SC, USA). To break cold dormancy, seeds were placed in sand-filled bags and kept at 4°C for two months prior to sowing. Approximately two months before the start of the experiment, seeds were sown into Lambert LM-GPS germination soil and placed in a temperature-controlled greenhouse room that was held between 25°C and 29.4°C. *A. incarnata* germination rates tend to be relatively low, so seed trays were topped with vermiculite to aid in moisture retention. Seedlings were fertilized with approximately 20 PPM of Jack’s LX 15-5-15 with 4% Ca and 2% Mg fertilizer three times a week until the majority of plants grew two sets of true leaves. All plants were then re-potted into Pro-mix BK25 soil, moved to a new temperature-controlled room that was held between 25.6°C and 29.4°C, and fertilized three times a week as described above. Approximately one week before the start of the experiment, plants were moved into the same greenhouse room that caterpillars were reared in (described below).

#### 1.3 *D. plexippus* Rearing

Monarch butterflies were caught and labeled near St. Marks, Florida, U.S.A. (30°09’33”N 84°12’26”W) between October 21st and October 23rd, 2022. Clear tape was placed on the abdomen of each butterfly and examined under a stereomicroscope to ensure they were not infected by *Ophryocystis elektroscirrha*, a common parasite of monarch butterflies. Prior to mating season, wild-caught monarch butterflies were stored in glassine envelopes at 14°C to induce a state of diapause, and were fed approximately 10-20% honey water every 10 days. Between March 6th and March 15th, 2023, wild-caught monarchs were placed in mesh cages for mating. Each cage was set up in a climate-controlled growth chamber (25°C, 16-hour/8-hour day/night cycle) and contained three male and three female butterflies. All cages were provided with a petri dish containing a sponge soaked in approximately 10-20% honey water for butterfly feeding. Mating cages were checked every 14 hours, and copulated butterflies were transferred to their own separate cage. After a copulated pair had detached the next day, the male was removed from the cage and the female was given a potted *A. curassavica* plant for oviposition, as well as honey water as described above. After a given female was done laying eggs, the plant was taken out of the growth chamber and placed in a temperature-controlled greenhouse room that was held between 23.3°C and 27.8°C for.

F1 caterpillars were reared on *A. curassavica* in the same greenhouse room previously described. After pupation, the silk attached to the end of the pupal cremaster was used to hot glue the pupae to the lid of clear solo cups, which were then taken from the greenhouse to the laboratory ( 22°C) for eclosion. A piece of paper towel was placed in the bottom of cups to help absorb liquids produced during the eclosion process. After eclosion, butterflies were placed in glassine envelopes and stored as previously described.

Between April 23rd and May 1st, 2023, F1 butterflies from different lineages and that were not infected with *O. elektroscirrha* were mated as previously described in the F0 generation. F1 females were given either *A. curassavica* or *A. incarnata* for oviposition, and caterpillars were collectively placed on their treatment 2 plant species upon hatching. Care was taken to make sure caterpillars that had taken bites of the plant they were oviposited on to were placed on the same milkweed species. Likewise, only caterpillars that had not taken any bites of the plant they were oviposited on were placed on the other milkweed species. To reach the sample size needed for this experiment, we used F2 caterpillars from two different lineages that did not share F0 or F1 ancestors. Treatments of plant species and development stage were randomly distributed to caterpillars from both lineages to minimize confounding due to genetic background.

#### 1.4 Sampling across life stages

To minimize changes in transcription due to sample handling, all caterpillars, pupae, and adults were snap frozen in liquid nitrogen before being stored in -80°C. Third instar caterpillars were pulled from their feeding plant and quickly placed into a sterile 2mL microcentrifuge tube that was then dipped in liquid nitrogen. Fifth instar caterpillars were frozen in the same way but were placed in sterile 5mL centrifuge tubes. Caterpillars that ate all of the leaves off of the plant they were originally placed on were placed on another plant of the same species.

One day after pupation, early pupae were placed in 5mL centrifuge tubes and frozen in liquid nitrogen as described above. Three days after pupation, pupae assigned to late pupa and adult stages were removed from their plant and taped to the lids of clear solo cups using silk attached to the cremaster. In some cases, not enough silk detached with the pupa, and tape was applied directly to the cremaster. Solo cups were then placed on the bottom rack of the same shelf that the caterpillars were reared on, and shade was provided by placing plastic trays above and to the southeast facing side of the shelf to prevent pupae from burning. A piece of paper towel was placed in the bottom of the cups containing adult samples to absorb fluids produced during the pupation process. Since there is variation in how long it takes for a pupa to eclose, late pupae were collected 6-8 days after pupation. Care was taken to ensure the distributions of how many days after pupation late pupae were sampled were equal between plants. Adults were frozen several hours after eclosion to allow their wings to fully expand. Here, adults were removed from their solo cup and quickly placed in glassine envelopes, which were then quickly frozen in liquid nitrogen.

After flash freezing in liquid nitrogen, samples were stored in a styrofoam cooler full of dry ice until all freezing for that day was completed. This process took approximately one hour or less on any given day, so no sample was on dry ice for more than an hour before being transferred to the -80°C freezer. All freezing took place in the same greenhouse room that the caterpillars were reared in, and no monarch left said room before being frozen throughout the duration of the experiment.

#### 1.5 RNA extraction and sequencing

We use a Promega SV Total Isolation System kit to extract total RNA from the monarch homogenate. Extractions were performed in batches of 11 samples with 1 negative control (per extraction batch). After each extraction, we used a NanoDrop to quantify the purity and concentration of the RNA. Samples with an A260/A280 or an A260/A230 of less than 1.95 were discarded and re-extracted to meet purity standards. While the general workflow followed the manufacturer’s suggested protocol, we made some alterations to obtain higher quality RNA extract. Briefly, we doubled the recommended RNA lysis buffer to decrease the tissue concentration in the initial lysis step. All centrifugation steps were increased to 20,000 rcf to better remove organic contaminants and performed at 17°C to avoid sample heating. We also added an additional centrifugation step after the initial tissue lysis to further clear organic contaminants and improve final extract purity. The specific protocol is as follows:

1. Add homogenate to 2mL microcentrifuge tube.
2. Immediately add 590 uL of RNA Lysis Buffer (RLA+BME) into microcentrifuge tube with homogenate
3. Use sterile micropestle to crush and lyse monarch homogenate (vigorously crush and spin pestle in tube for approximately 1 minute).
4. Centrifuge at 20,000 rcf for 10 minutes at 17°C.
5. Transfer approximately 400 uL to 500 uL of aqueous phase to a new microcentrifuge tube.
6. Centrifuge at 20,000 rcf for 20 minutes at 17°C.
7. Transfer 175uL of the cleared lysate (aqueous layer) to a new microcentrifuge tube.
8. Add 200 uL of 95
9. Transfer lysate+ethanol to Spin Basket Assembly and centrifuge for 5 minutes at 20,000 rcf and 17°C.
10. While the centrifuge is running, Prepare DNase incubation mix: 40uL of Yellow Core Buffer + 5uL MnCl2 + 5uL of Dnase I (per sample). Mix gently via pipetting.
11. Discard eluate.
12. Add 50 uL of DNase incubation mix to the membrane of the Spin Basket, incubate for 15 minutes at room temperature.
13. Add 200 uL of DNase Stop Solution (DSA+ethanol) and centrifuge at 20,000 rcf for 1 minute at 17°C.
14. Discard eluate.
15. Add 600 uL of RNA Wash Solution (RWA); centrifuge at 20,000 rcf for 1 minute at 17°C.
16. Discard eluate.
17. Add 250 uL of RNA Was Solution (RWA); centrifuge at 20,000 rcf for 2 minutes at 17°C.
18. Transfer Spin Basket to Elution Tube.
19. Add 100 uL of Nuclease-Free water to the Spin Basket membrane.
20. Centrifuge at 20,000 rcf for 1 minute to elute RNA.
21. Store at -80°C.

After all extractions were completed, purified RNA was packaged in dry ice and sent to Novogene for sequencing. Briefly, Novogene used an Agilent 5400 Fragment Analyzer System to performed additional quality control. This involved reconfirming sample purity, ensuring that all samples had adequate concentrations and volumes, and checking that all sampled had acceptable RNA integrity numbers (minimum = 7.9). After additional quality assessment, mRNA was separated via poly-A tail selection, and 150bp paired-end sequencing was performed using a NovaSeq 6000 sequencing system, ensuring at least 20 million reads were obtained for each sample.

#### 1.6 Sequence processing and gene expression quantification

Quality control of raw sequences was initially performed by Novogene. This entailed the removal adapter sequences, the removal of reads with ambiguous base calls in greater than 10% of the read, and the removal of reads with a phred score of less than or equal to 5 in 50% of the read. After receiving the sequences from Novogene, we used FASTQC to generate additional quality reports for each sample [2]. This showed that for each sample, the median phred score did not drop below 30 at any position along the reads. Therefore, no additional quality control was performed.

To quantify transcript abundances for each gene, we used kallisto (v.0.46.2) to pseudo-align reads to the coding sequences of the D. plexippus reference genome (v.Dpv3; GCA 000235995.2) [3]. Downstream analyses were performed using transcript per million normalized read counts (automatically generated by kallisto) to minimize biases due to unequal gene lengths and varying library sizes [4, 5].

#### 1.7 Quantifying gene expression divergence between stages

Given the high dimensionality of gene expression data, we first computed the Manhattan distance between each sample using the dist R function [6]. We then used the adonis2 function from the vegan R package (v.2.6-4) [7] to perform a permutational multivariate analysis of variance (PERMANOVA) with 999 permutations, where developmental stage and plant were initially considered as factors. We then performed a PERMANOVA on each set of adjacent stages, as well as between each larval stage and the adult stage. To visualize global expression divergence between stages, we performed principal component analysis using the *prcomp* R function [6].

#### 1.8 Inferring homologous gene groups

To infer homology between genes, we first used PSI-BLAST (BLAST 2.5.0+) [8] with five iterations to align all D. plexippus protein sequences to each other. Genes were then inferred to be homologous if the query sequence showed at least 30% similarity across the length of the target sequence, as well as an E-value of at least 1x10-10. To examine how including more distant homologs could impact our analysis, we performed an additional analysis where homology was inferred based on at least 20% similarity across 70% of the target sequence and an E-value of less than 1x10-5. Homologous pairs were assembled into sets of two-node subgraphs, and subgraphs were then merged based on common node identity to assemble homologous groups. To quantify the phylogenetic distance between members of inferred homologous gene groups, we first used MUSCLE (v.5.1) to create a multiple sequence alignment for each group [9]. We then used IQ-TREE2 (v.2.1.4) to identify the best fit sequence evolution model and infer maximum likelihood phylogenies for each multiple sequence alignment [10, 11].

#### 1.9 Quantifying the relationship between gene phylogenetic divergence and expression pattern divergence within homologous groups

To quantify the relatinship between phylogenetic divergence and expression divergence within homologous groups, we used the *cophenetic.phylo* function from the *ape* R package (v. 5.7-1) [12] to calculate pairwise phylogenetic distances from each homologous group tree. To calculate pairwise expression pattern distances, we first mean centered and standardized the median transcripts/million value for each gene within each stage to better measure distance between temporal patterns as opposed to magnitude (which cannot be assessed with our data). We then calculated the pairwise Euclidian distance between each gene expression pattern within a given homologous group using the *dist* R function [6]. Finally, we used Mantel tests to calculate the correlation between phylogenetic and expression pattern distance matrices for each homologous group, which were implemented via the *mantel* function in the vegan R package (v.2.6-4) [7]. We then used a t-test to test if the distribution of correlation coefficients was positively shifted from 0, which was implemented using the *t.test* R function [6].

#### 1.10 Quantifying the relationship between phylogenetic and expression pattern diversity

The diversity (D) of each tree was then calculated by summing all branch lengths: *D* = *^n^ l_i_*, where *n* represents the number of branches and *l_i_* represents the length of the *ith* branch. To quantify expression pattern diversity, we first created hierarchical clustering graphs of the temporal expression patterns for each gene using the Ward method, as implemented by hclust R function [6]. Prior to clustering, the transcripts/million values for each gene were mean centered and standardized because hierarchical clustering will group expression patterns that show distinct temporal trends but have more similar relative abundances at each time point. For each hierarchical clustering graph, diversity was calculated as previously described for phylogenetic diversity. We then fit a linear model to examine the relationship between phylogenetic diversity and expression pattern diversity across all inferred homologous gene groups, which was implemented using the *lm* R function [6]. Because diversity was calculated additively (for each branch, diversity was added in proportion to divergence), we also fit individual linear models to each homologous gene group size that had at least five replicates. In addition to removing the inherent positive correlation between group size and diversity, this approach also allowed us to contrast global and local patterns.

#### 1.11 Null model algorithm and analysis

To collect stronger evidence regarding the relationship between phylogenetic and expression pattern diversity, we wanted to test if the observed patterns could be randomly derived from our data. Therefore, we constructed a null model that assigns a randomly sampled expression pattern to each gene for each homologous gene group and recalculates the linear relationship between phylogenetic and expression pattern diversity.

Consider the total pool of observed expression patterns *E* = *{X*_1_*, X*_2_*, X*_3_*, . . ., X_i_}*, where *X* represents an expression pattern and *i* represents the number of genes in the *D. plexippus* genome. Previously defined groups of homologous genes are represented as *H* = *{G*1*, G*2*, G*3*, . . ., G_j_}*, where *j* represents the number of homologous gene groups and also serves as the index for the set of observed phylogenetic diversities for each group, represented as *P* = *{D*_1_*, D*_2_*, D*_3_*, . . ., D_j_}*. For each group *G_j_*, *Gj* = *{X_i_*_1_ *, X_i_*_2_ *, X_i_*_3_ *, . . ., X_i__n_ }*, where *n* represents the number of genes included in each group and *X_i__n_* represents an expression pattern from *E*. Note that not all expression patterns in *E* are included in some *G_j_*. For each *X_i__n_* in each *G_j_*, *X_i__n_* ← *X_i__k_*, where *X_i__k_* represents the *kth* expression pattern randomly selected with replacement from *E* for the *jth* group. After random assignments of expression patterns, expression pattern diversity *S_j_* is calculated for each *G_j_* as previously described, which is represented as *R* = *{S*_1_*, S*_2_*, S*_3_*, . . ., S_j_}*. *R* is then combined with *P* to produce *N* such that *N* = *{*(*D_j_, S_j_*)*|D_j_ ∈ P, Sj ∈ R}*. Finally, a linear model is then fit both globally and to each group size, as previously described for the observed analysis.

This algorithm was repeated 1000 times to generate null distributions of global and group-size specific relationships between phylogenetic and expression pattern diversity. To evaluate the observed results against the null model predictions, we calculated the probability that the null model would produce an value greater than the observed, as well as the standardized effect sizes 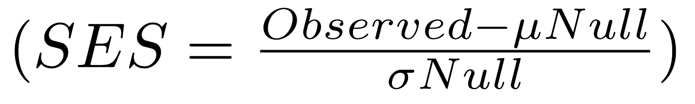 for the slope and explained variance (*R*^2^) of each linear model.

#### 1.12 Expression specificity calculation and analysis

Stage-specificity for each gene was calculated using the tissue specificity index *τ* [13], which ranges from 0 (broad expression) to 1 (specific expression): 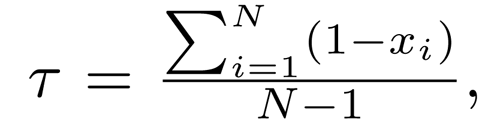 where *N* is the number of stages (for our *N −*1 purposes) and *x_i_* is the expression level normalized to the maximum expression value across stages. Although *τ* was developed for assessing tissue specificity, it has been used to gain insight into temporal specificity as well [14]. We then performed a Kolmogorov–Smirnov test using the *ks.test* R function [6] to assess if the distribution of *τ* values was shifted in duplicated genes relative to singleton genes.

### 2 Methodological Summaries

#### 2.1 RNA quality control report

**Table 1:** RNA extract quality control report.

| Sample Name | Concentration (ng/ul) | Volume (ul) | Total amount (ug) | RIN |
| --- | --- | --- | --- | --- |
| mtstp3cu2 | 120.15 | 91 | 10.93 | 9.7 |
| mtstp3cu3 | 435.84 | 93 | 40.53 | 9.6 |
| mtstp3cu4 | 99.07 | 88 | 8.72 | 9.6 |
| mtstp3cu5 | 223.96 | 94 | 21.05 | 9.7 |
| mtstp3cu8 | 111.95 | 91 | 10.19 | 9.6 |
| mtstp3iu81 | 211.09 | 92 | 19.42 | 9.8 |
| mtstp3iu82 | 194.64 | 93 | 18.1 | 9.8 |
| mtstp3iu83 | 373.28 | 91 | 33.97 | 9.6 |
| mtstp3iu84 | 196.04 | 93 | 18.23 | 9.6 |
| mtstp3iu85 | 136.84 | 90 | 12.32 | 9.7 |
| mtstp5cu17 | 363.74 | 101 | 36.74 | 9.4 |
| mtstp5cu18 | 236.69 | 92 | 21.78 | 9.8 |
| mtstp5cu19 | 544.95 | 93 | 50.68 | 9.5 |
| mtstp5cu20 | 130 | 91 | 11.83 | 9.7 |
| mtstp5cu21 | 400.58 | 94 | 37.65 | 9.8 |
| mtstp5iu100 | 377.26 | 91 | 34.33 | 9.7 |
| mtstp5iu101 | 98.09 | 89 | 8.73 | 9.6 |
| mtstp5iu97 | 164.64 | 92 | 15.15 | 9.8 |
| mtstp5iu98 | 258.2 | 91 | 23.5 | 9.8 |
| mtstp5iu99 | 139.84 | 91 | 12.73 | 9.8 |
| mtstpAcu65 | 132.79 | 92 | 12.22 | 9.5 |
| mtstpAcu66 | 106.76 | 90 | 9.61 | 9.5 |
| mtstpAcu67 | 37.05 | 94 | 3.48 | 8.9 |
| mtstpAcu68 | 94.26 | 91 | 8.58 | 9.4 |
| mtstpAcu69 | 176.26 | 91 | 16.04 | 9.8 |
| mtstpAiu145 | 162.84 | 92 | 14.98 | 9.6 |
| mtstpAiu146 | 283.45 | 92 | 26.08 | 9.6 |
| mtstpAiu147 | 51.58 | 92 | 4.75 | 9.2 |
| mtstpAiu148 | 89.37 | 92 | 8.22 | 9.3 |
| mtstpAiu149 | 307.63 | 91 | 27.99 | 9.7 |
| mtstpEcu33 | 203.48 | 92 | 18.72 | 7.4 |
| mtstpEcu34 | 244.57 | 91 | 22.26 | 8.7 |
| mtstpEcu35 | 491.84 | 89 | 43.77 | 7.9 |
| mtstpEcu36 | 220.03 | 92 | 20.24 | 8.6 |
| mtstpEcu38 | 220.92 | 93 | 20.55 | 8.6 |
| mtstpEiu113 | 333.32 | 93 | 31 | 8.9 |
| mtstpEiu114 | 446.98 | 91 | 40.68 | 7.9 |
| mtstpEiu115 | 182.76 | 92 | 16.81 | 8.6 |
| mtstpEiu116 | 257.7 | 89 | 22.94 | 8.4 |
| mtstpEiu117 | 326.41 | 92 | 30.03 | 8 |
| mtstpLcu49 | 95.18 | 92 | 8.76 | 9 |
| mtstpLcu50 | 93.51 | 92 | 8.6 | 9.3 |
| mtstpLcu52 | 220.52 | 93 | 20.51 | 8.8 |
| mtstpLcu53 | 95.67 | 91 | 8.71 | 9.4 |
| mtstpLcu56 | 399.94 | 89 | 35.59 | 8.2 |
| mtstpLiu129 | 145.89 | 91 | 13.28 | 9.1 |
| mtstpLiu130 | 91.59 | 91 | 8.33 | 8.7 |
| mtstpLiu131 | 391.6 | 87 | 34.07 | 9.6 |
| mtstpLiu133 | 85.42 | 89 | 7.6 | 8.4 |
| mtstpLiu135 | 108.7 | 92 | 10 | 8.1 |

#### 2.2 RNA sequencing statistics

After quantifying transcript counts per gene, we checked that our sequencing effort was adequate to down-stream analyses. First, we examined the number and proportion of raw reads that passed quality control, as well as the number and proportion of quality-controlled reads that were pseudo-aligned to the *D. plexippus* genome (Table 2). We then generated a rarefaction plot see if our sequencing depth had sufficiently detected the expression of most transcripts that were expressed at a given stage (Figure 1).

**Table 2:**
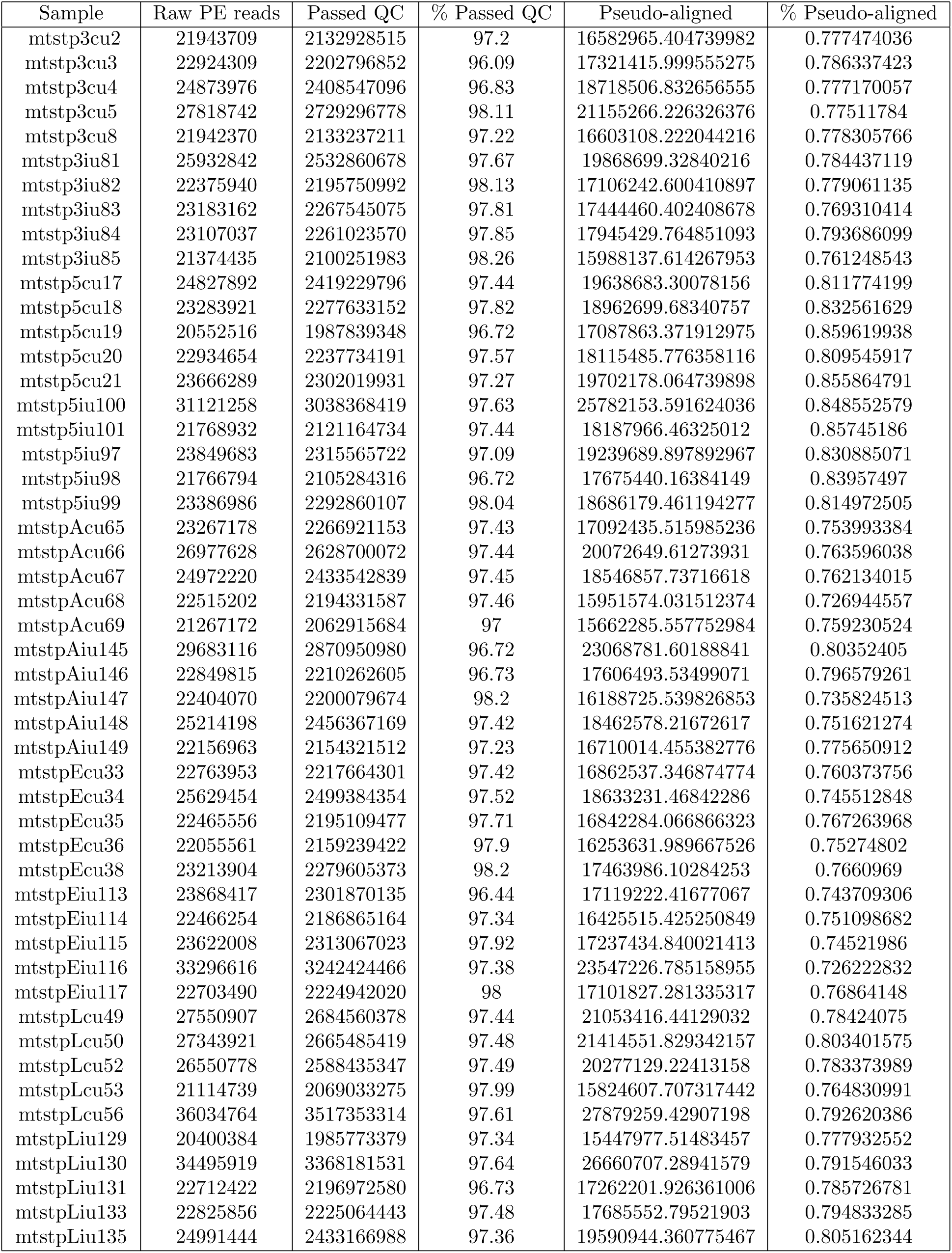
Sequence processing and mapping summary.

**Figure 1:**
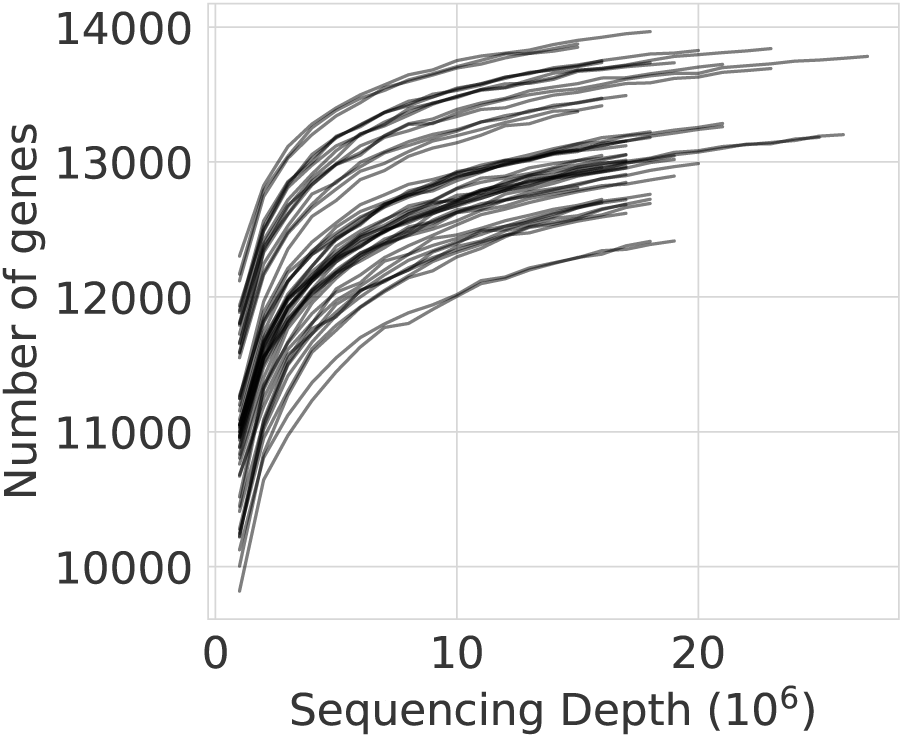
Rarefaction curves showing the number of genes detected on the y-axis and sequencing depth on the x-axis. Each line corresponds to an individual sample. Plateaus in the number of detected genes at higher sequencing depths suggest that our sequencing effort was sufficient.

#### 2.3 Summary of homologous gene group inferences

Homologous gene group inference is described in section 1.8. The following plots show the summary histograms of homologous group size.

**Figure 2:**
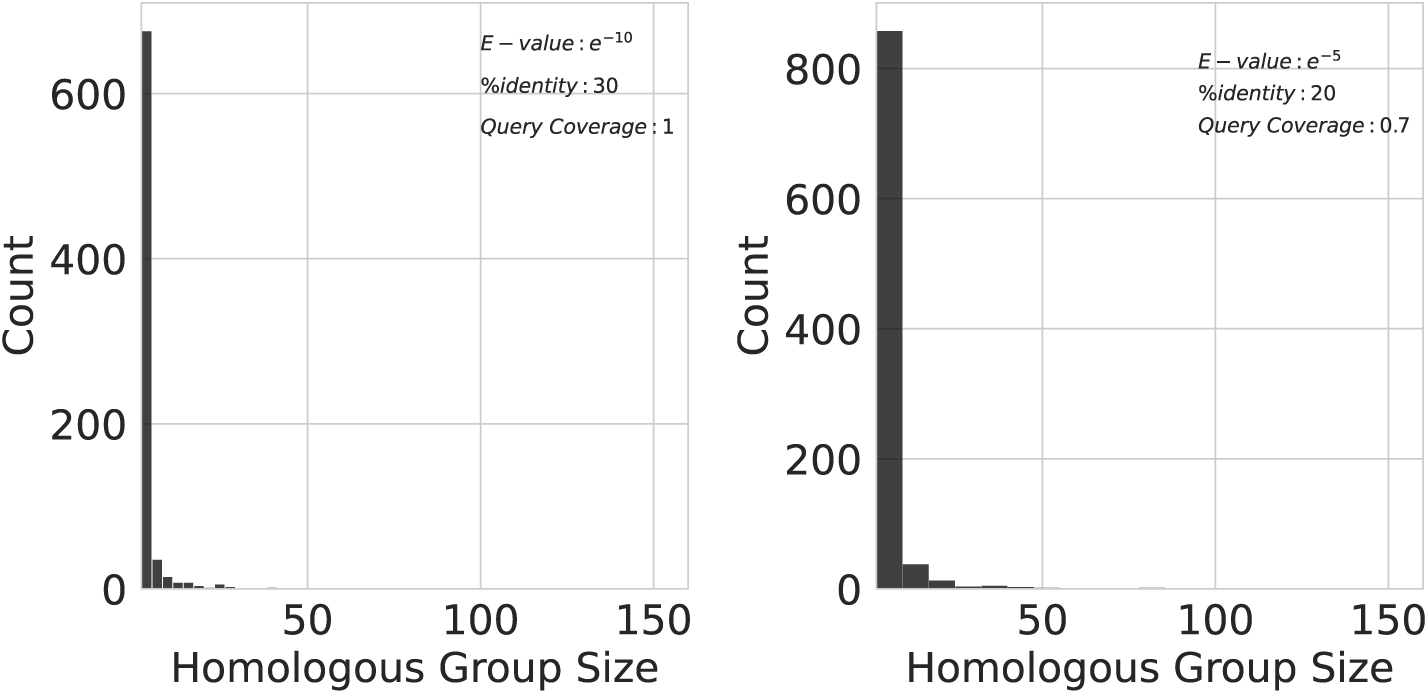
Histograms showing the distribution of homologous group sizes detected in the *D. plexippus* genome when using more (left) and less (right) stringent sequence similarity cutoffs.

### 3 Supporting Results

#### 3.1 Observed - Null model comparisons

To compare the observed slope(s) of the relationship between phylogenetic and expression pattern diversity to our null mode (described in 1.11), we calculated the standardized effect size and probability that the null model would generate an effect greater than or equal to the observed. Direct comparisons between observed and null relationships are depicted in Figure 3. The results for our analysis that used a more stringent criteria for inferring homology are reported in Table 3 and summarized in the main text.

**Figure 3:**
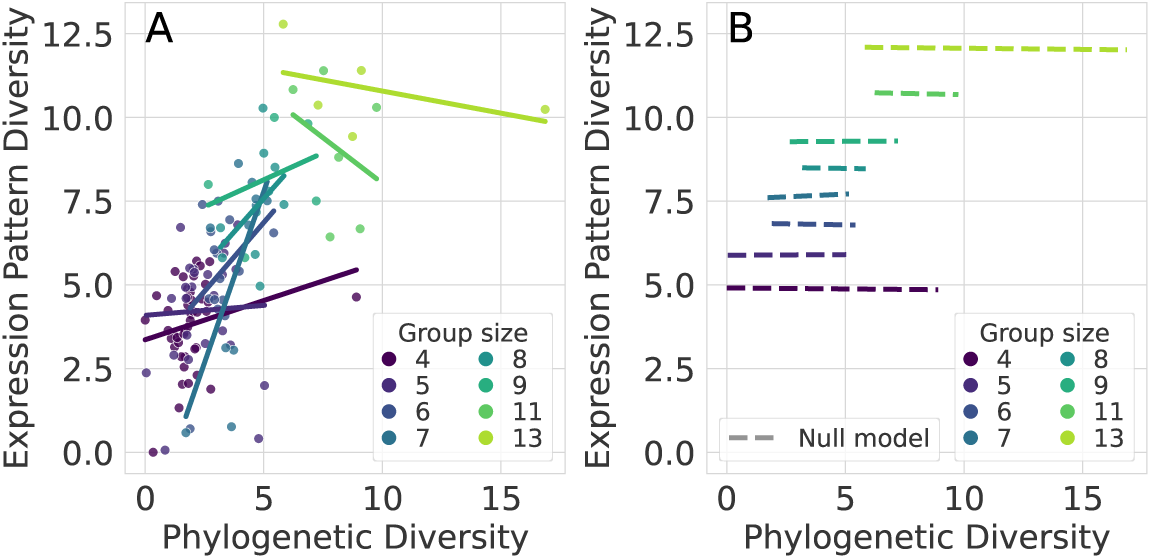
The relationship between phylogenetic and expression pattern diversity A) within each homologous gene group size, and B) as predicted by a null model that randomly assigns expression patterns to genes. A corresponds to Figure 4D in the main text and is placed here for direct visual comparisons to the null model predictions. In A, each line represents the linear model fit to each homologous group size (only group sizes will 5 or more replicates were considered in this analysis). In B, dashed lines represent the average null linear prediction for each homologous group size. Each line corresponds to an observed linear model depicted in A.

**Table 3:** A table showing the results for each observed-null model comparison that were summarized in the main text.

| Homologous group size | Standardized effect size (SES) | $P(Null \geq Observed)$ |
| --- | --- | --- |
| Global | -4.075340 | 1 |
| 4 | 1.999710 | 0.014 |
| 5 | 0.348024 | 0.351 |
| 6 | 2.444219 | 0.009 |
| 7 | 6.110705 | 0.000 |
| 8 | 1.990671 | 0.025 |
| 9 | 1.152409 | 0.131 |
| 11 | -1.298748 | 0.908 |
| 13 | -0.893563 | 0.810 |

#### 3.2 Reanalysis based on less stringent homology inference

To ensure that our findings were robust, we re-analyzed our data using less stringent sequence identity cutoffs to infer gene homology (see section 1.8). This included more divergent genes in our analysis, thus increasing the amount of phylogenetic diversity captured.

##### 3.2.1 Correlations between phylogenetic and expression pattern diversity

The overall relationships between phylogenetic and expression pattern diversity were consistent with our primary analysis (Figure 3). The only notable difference is that the expression pattern diversity increases at a lower rate with respect to phylogenetic diversity (slope = 0.64, SES = -5.47, *P* (*Observed < Null*) *<* 0.001) (Figure 3A). This is expected given the findings presented in the main text, where the rate at which expression diversity increases with phylogenetic diversity declines at more phylogenetically diverse groups. Furthermore, inspection of global-null comparisons are also consistent with the findings presented in the main text, where the deviation from null expectations deteriorates at larger and more diverse groups.

##### 3.2.2 Expression pattern specificity

Our comparisons of stage-specificity between duplicate and singleton genes based on less stringent similarity cutoffs for homology inference were consistent with the analysis presented in the main text. Specifically, genes that are part of homologous groups tend to show increased stage-specificity relative to singleton genes (D = 0.10, *p <* 2.2 *∗* 10*^−^*^16^). Although this pattern is statistically supported, we note that the effect size is slightly smaller than the analysis presented in the main text analysis.

**Figure 4:**
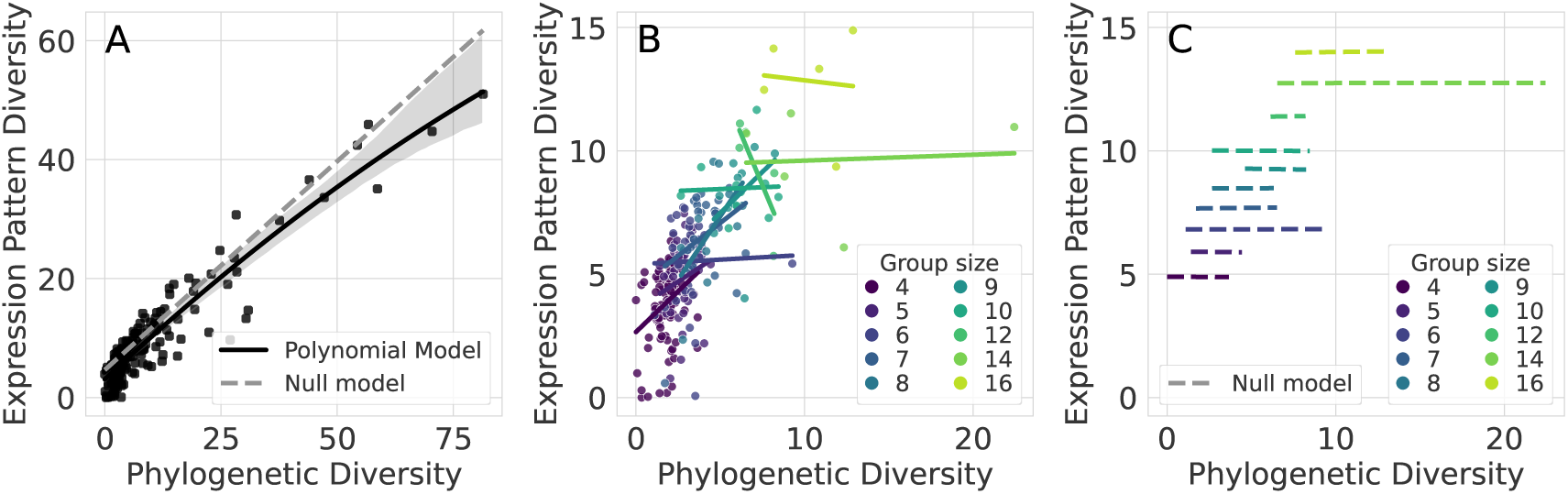
The relationship between phylogenetic and expression pattern diversity A) across all homologous gene groups, B) within each homologous gene group size, and C) as predicted by a null model that randomly assigns expression patterns to genes. In A, the solid black line depicts the fit polynomial model, the light gray area indicates the 95% confidence interval for said model, and the dashed line shows the average null linear prediction. In B, each line represents the linear model fit to each homologous group size (only group sizes will 5 or more replicates were considered in this analysis). In C, dashed lines represent the average null linear prediction for each homologous group size. Each line corresponds to an observed linear model depicted in B.

**Table 4:** A table showing the results for each observed-null model comparison that were produced by our reanalysis based on less stringent similarity cutoffs for homology inference.

| Homologous group size | Standardized effect size (SES) | $P(Null \geq Observed)$ |
| --- | --- | --- |
| Global | -5.472289 | 1 |
| 4 | 4.794734 | 0.000 |
| 5 | 2.217783 | 0.018 |
| 6 | 0.262302 | 0.418 |
| 7 | 2.597285 | 0.005 |
| 8 | 3.667600 | 0.000 |
| 9 | 1.892611 | 0.026 |
| 10 | 0.149418 | 0.440 |
| 12 | -2.954615 | 0.999 |
| 14 | 0.231405 | 0.413 |
| 16 | -0.334492 | 0.629 |

**Figure 5:**
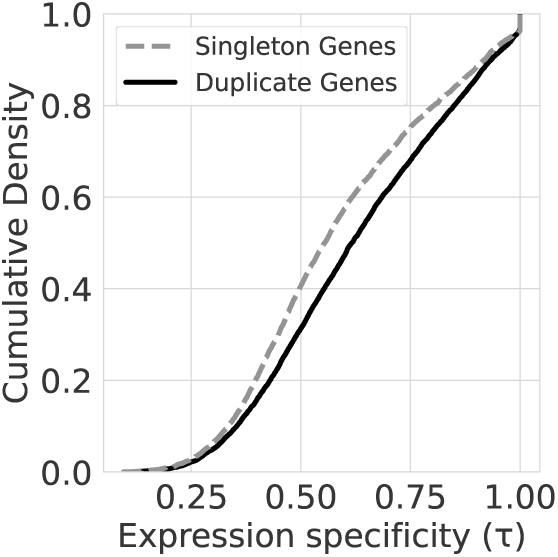
The empirical cumulative density functions of expression specificity values *τ* for duplicate (solid line) and singleton (dashed line) genes. Higher expression specificity values indicate more stage-specific expression patterns.

#### 3.3 Broad functional overview

To gain a general sense of what high level functional differences occurred between stages, we used the KEGG [15] to infer gene functions and examined the relative transcriptional investment in the highest level KEGG BRITE groupings (Figure 6).

**Figure 6:**
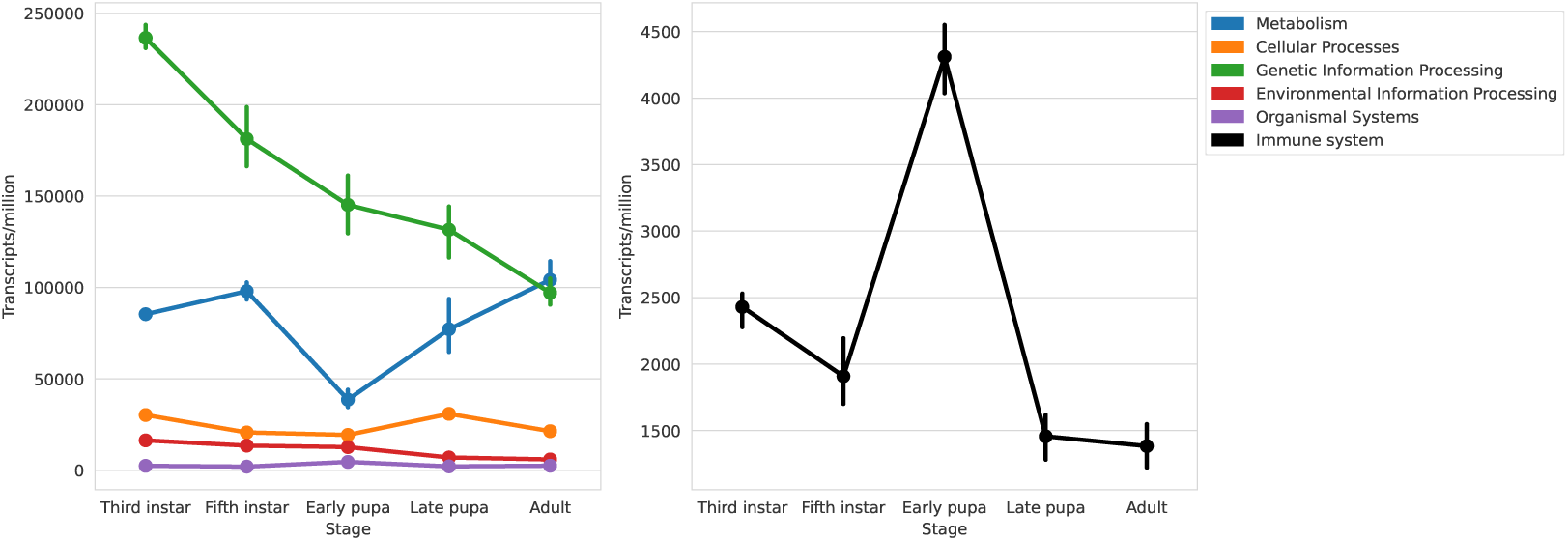
A line plot showing the relative transcriptional investment (transcripts per million) in each high level functional group across life stages. Note that these groupings reflect the overall transcriptional investment in each broad functional group listed, not the activity of individual pathways or genes. Therefore, each individual pathway or gene within each group is not expected to necessarily follow the exact trend exhibited by the whole group. Error bars represent 95% confidence intervals calculated across individual samples.

